# Architecture of a portal complex embedded in the poxvirus core

**DOI:** 10.64898/2026.06.09.730874

**Authors:** Ye Hong, Shiyuan Liu, Rui Liang, Keren Wang, Junxia Zhang, Cheng Peng, Zefeng Zhu, Zheyuan Zhang, Weiping Zhang, Baoying Huang, Chen Song, Wenjie Tan, Sai Li

## Abstract

Gene transcription within the viral core is a unique feature of poxvirus replication. Following entry into the cytoplasm, the poxvirus core dissociates from the lateral bodies and undergoes expansion, functioning as a compartment for early transcription. However, the mechanisms governing molecular exchange between the viral core and the host cytoplasm remain poorly understood. Here, we determine the structures of the portal complex and its pore on the poxvirus core at 7.1 Å and 4.9 Å resolution, respectively, using cryo-electron tomography and sub-tomogram averaging. The pore is assembled from three viral proteins, E8, E6, and L3, for which we constructed an atomic model. Structural and channel analyses reveal that the pore satisfies the geometric and electrostatic requirements for the transport of RNA and smaller negatively charged molecules, while excluding double-stranded DNA and cytosolic DNA sensors. Together, our findings establish a structural framework for understanding the assembly and function of the poxvirus portal complex and identify potential targets for antiviral intervention.

## Introduction

The genus Orthopoxvirus (OPXV), within the family *Poxviridae* and subfamily *Chordopoxvirinae*, comprises large double-stranded DNA viruses, many of which pose substantial threats to human health. Prominent members include variola virus (VARV), the causative agent of smallpox; and monkeypox virus (MPXV), the etiological agent of mpox. In recent years, recurrent mpox outbreaks have twice been designated as Public Health Emergencies of International Concern by the World Health Organization (WHO), underscoring the persistent global risk posed by OPXVs. Among these viruses, vaccinia virus (VACV) serves as the most extensively studied prototype. VACV contains an approximately 200 kbp linear double-stranded DNA genome encoding nearly 200 proteins and shares greater than 90% nucleotide identity with both VARV and MPXV. Importantly, VACV has been widely engineered as a vaccine platform conferring protection against smallpox and mpox.

VACV exists in two infectious forms: the mature virus (MV) and the enveloped virus (EV). The MV comprises a biconcave viral core and two lateral bodies enclosed within a viral envelope, whereas the EV acquires an additional outer membrane. During infection, the EV outer membrane ruptures, delivering the underlying MV into host cells. Recently, we and other groups have refined the architectural understanding of OPXVs, revealing *in-situ* structures and spatial arrangements of several core-associated proteins^1–7^. The viral core, the defining structural compartment of MVs and EVs, encases the viral genomic DNA within a proteinaceous wall composed of two layers. The outer palisade layer is formed by trimeric p4a (the processed and cleaved form of A10) together with A4 (also termed p39)^1–3^, while the inner wall layer consists predominantly of continuous p4b (processed and cleaved A3)^1^. Occasionally, portal complexes composed of six palisade trimers arranged around a pore are observed breaching the core wall^1–3,5–7^, forming the largest discernible openings on the viral core.

Following membrane fusion and cytoplasmic entry, the VACV core undergoes a pronounced morphological transition from a biconcave to a brick-shaped architecture and initiates early transcription within its interior without core wall rupture^8^. This process necessitates highly selective molecular transport across the core wall: ribonucleotides must be imported, viral mRNA exported, and the genomic DNA shielded from cytosolic DNA sensors. Structural studies indicate that both the palisade and inner wall layers are densely packed, leaving limited interstitial space insufficient for the passage of large macromolecules. The sporadic observation of pores penetrating the core wall has therefore suggested the existence of specialized conduits mediating molecular exchange. However, the precise protein composition and structural details of these pores have remained unresolved.

Here, we present the structure of the VACV portal complex at 7.1 Å resolution and its pore at 4.9 Å resolution, determined from isolated viral cores using cryo-electron tomography (cryo-ET) and sub-tomogram averaging (STA). Guided by structural prediction and prior functional studies, we identify E8, E6, and L3 as the principal components of the pore. Within the central channel of the pore, the narrowest constriction is lined by four lysine residues, each of which is essential for virus replication. These findings provide mechanistic insight into portal complex architecture and establish a structural basis for understanding early transcription in poxviruses.

## Results

### Architecture and distribution of portal complexes on the naked VACV core

Natively released cores observed *in situ* adopt a brick-shaped morphology enclosed by both palisade and inner wall layers (Fig. 1a). Flower-shaped portal complexes were occasionally detected on the top and bottom surfaces of the cores (Fig. 1b). To facilitate structural characterization, we isolated MVs from infected cells and treated them with 1% NP40 and 1 mM dithiothreitol (DTT) at 37 °C to remove the viral envelope, thereby releasing viral cores. This treatment has previously been shown to yield morphologically intact cores^3^ that retain transcriptional activity upon supplementation with ribonucleotides^9,10^. Cryo-ET confirmed that purified cores preserved their brick-shaped architecture and remained densely coated with palisade trimers. Distinct portal complexes were readily observed on the flat top and bottom surfaces (Fig. 1c-d). Notably, neither the global core architecture nor the structural features of the portal complexes differed between purified cores and cores naturally released within host cells.

**Fig. 1.**
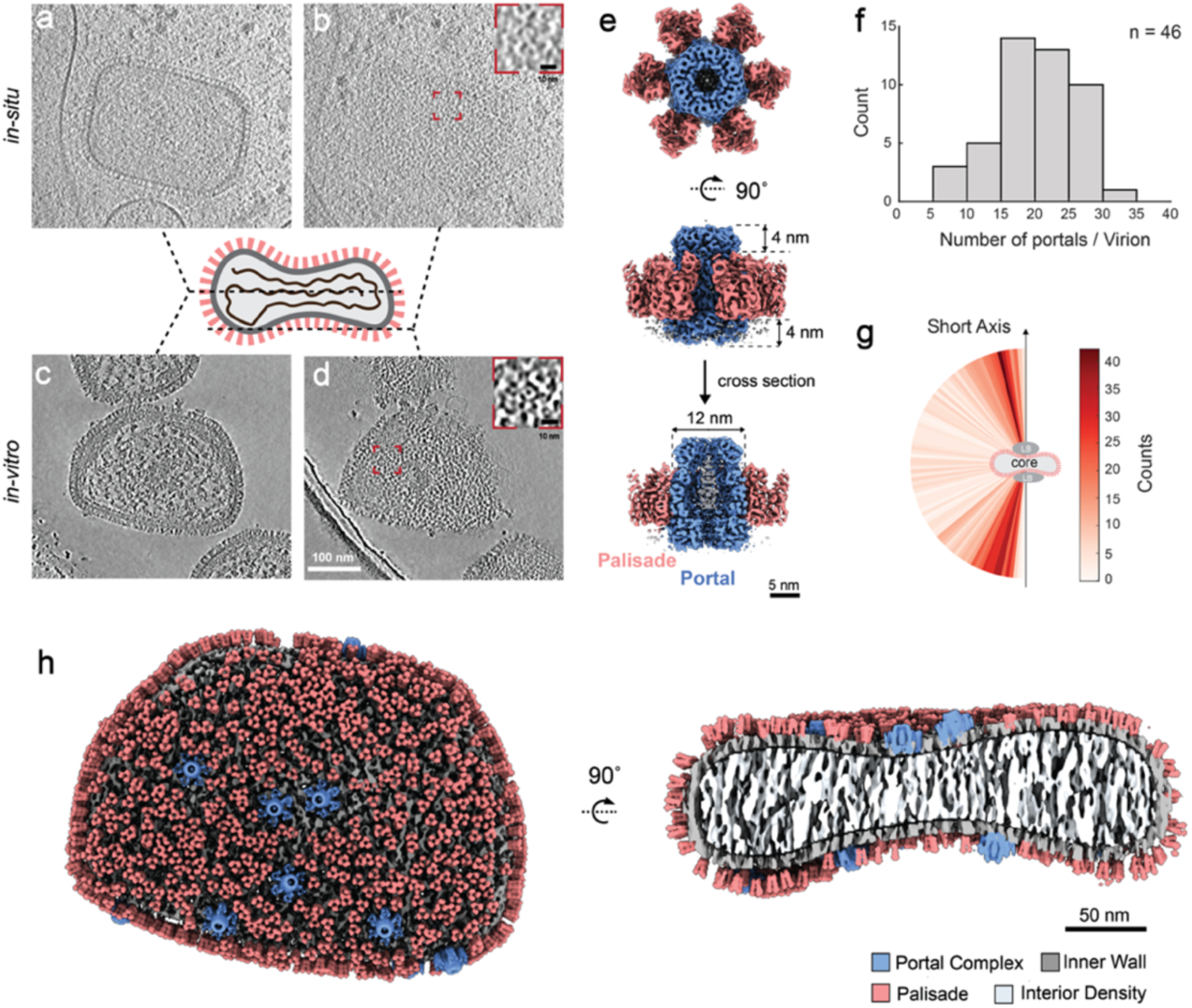
Architecture and distribution of the portal complexes on VACV core. **a-b**, Tomogram slices showing the central slice (**a**) and bottom surface (**b**) of an exemplary natively released core in cytosol imaged at the cell periphery. Thickness of the tomogram slice is 16 nm. **c-d**, Tomogram slices showing the central slice (**c**) and bottom surface (**d**) of an exemplary purified VACV core. Thickness of the tomogram slice is 5 nm. Inset red box on the top right of **b** and **d** show the zoom-in view of the portal complex. **e**, Top, side and cut-open views of the structure of the portal complex, comprising six palisades (salmon) surrounding the pore (blue) and a central plug density (grey). **f**, Histogram showing the number of portal complex on each virus core. **g**, portal complex orientation relative to the short axis of the virus core. 46 intactly imaged cores and 904 portal complexes were used for the statistics in (**f**) and (**g**). **h**, Top view and cut-open side view of a composite structure of the VACV core in **c** reconstructed by projecting the palisade and portal complex structures onto the densities of inner wall and core interior segmented from the tomogram. Only particles with high confidence were displayed in the composite structure. Misaligned particles were removed based on geometric constraints and cross-correlation.

Owing to the improved contrast of purified samples, we determined the structure of the portal complex at 7.1 Å resolution using cryo-ET and STA (Fig. 1e; Supplementary Figs. 1-2; Supplementary Table 1). The reconstruction revealed six surrounding palisade trimers arranged around a cylindrical pore, forming a hexamerically symmetric assembly. The pore, approximately 12 nm in diameter, protrudes ∼4 nm above and extends ∼4 nm below the palisade layer, traversing both layers of the core wall and thereby connecting the interior and exterior of the viral core. A continuous channel runs through the pore, with a non-occluding plug-like density visible at its center (Fig. 1e).

Local resolution analysis indicated that the pore region achieved higher resolution than the surrounding palisades and the central plug density, likely reflecting greater structural homogeneity (Supplementary Fig. 2b). We next examined the abundance and spatial distribution of portal complexes. On average, each VACV core contained 19.7 ± 5.6 portal complexes (Fig. 1f). Although portal complexes were occasionally observed near core edges, the majority localized to the flat surfaces and were oriented approximately parallel to the short axis of the core (Fig. 1g,h). Angular distribution analysis and structural features suggest that the final reconstruction is not significantly affected by preferred orientation (Supplementary Fig. 1b,c). No evident spatial pattern was observed in the distribution of portal complexes across the core surfaces (Fig. 1h).

### Sub-tomogram averaging resolves the pore structure at 4.9 Å resolution

To further improve structural resolution of the pore, we applied a focused mask encompassing only the pore region. This approach yielded a reconstruction at 4.9 Å resolution (Fig. 2a,b; Supplementary Fig. 3), enabling detailed structural dissection. The pore architecture comprises four concentric hexameric rings organized into three distinct layers (Fig. 2c). These include a “collar” ring at the top (Fig. 2d), a “coat” ring forming the central scaffold (Fig. 2e), and two rings at the base: an outer “skirt” ring (Fig. 2f) and an inner “petticoat” ring (Fig. 2g).

**Fig. 2.**
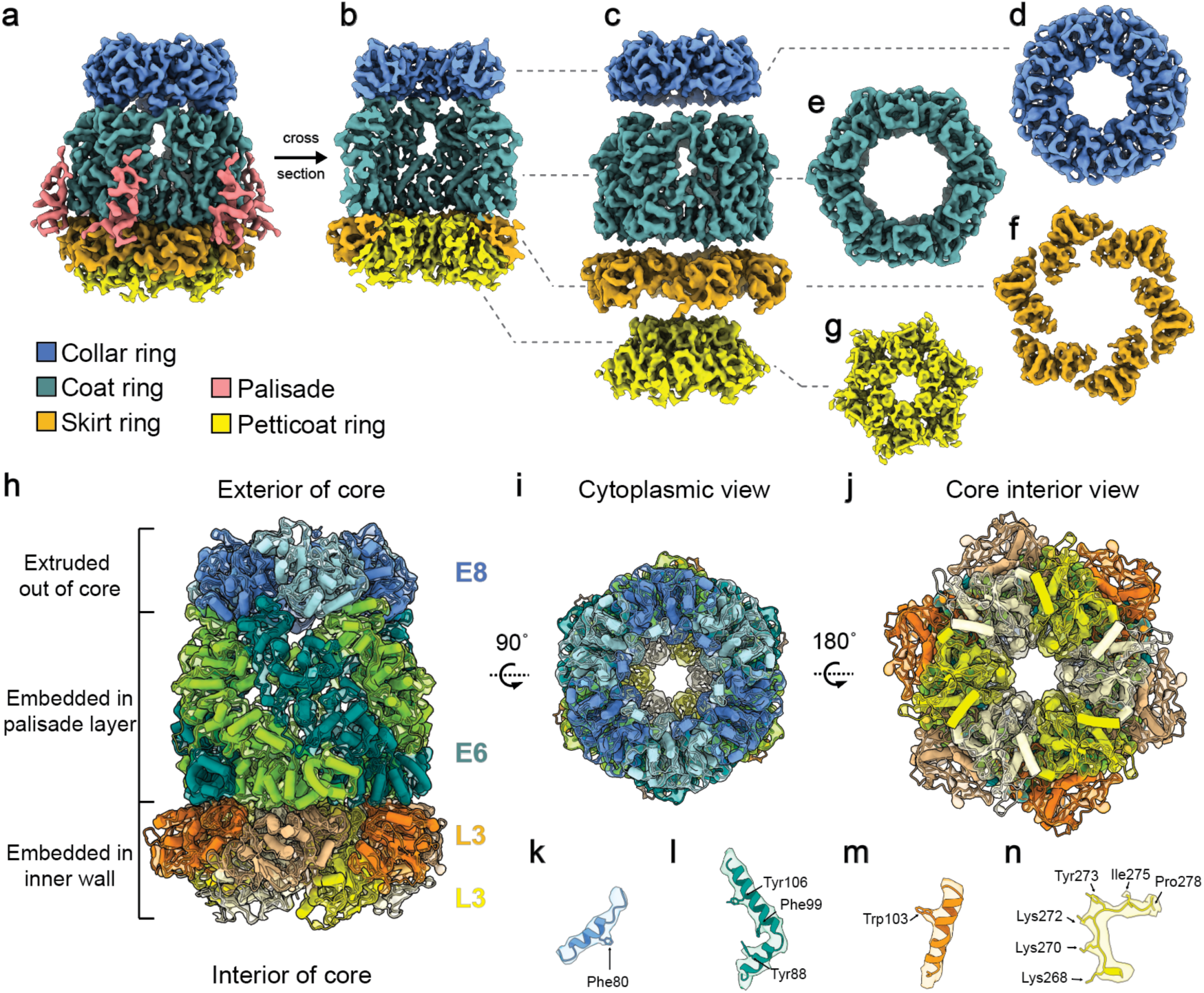
Structural dissection of the pore. **a**, The side view and **b**, cut-open view of the pore structure. **c**, The four rings composing the pore include the collar-ring at the top layer (blue), the coat-ring at the middle layer (green), and a bottom layer comprising an outer skirt-ring (orange) and an inner petticoat-ring (yellow). The palisade densities connecting to the pore in **a** are in salmon. **d-g**, Top view of the collar-, coat-, skirt- and petticoat-rings. **h-j**, The segmented density map of the pore fitted with the E8-E6-L3 complex model (E8/E6/L3=6:6:12) presented in three views. **k-n**, The zoom-in view of density on the collar-, coat-, skirt- and petticoat-rings fitted with their corresponding protein models.

The coat-ring constitutes the principal structural element of the pore. It is embedded within the palisade layer and mediates connectivity between the collar and skirt-rings. Among these structural components, the petticoat-ring forms the narrowest region of the channel, whereas the coat-ring defines the widest opening. At this resolution, secondary structure elements and bulky side chains were clearly discernible in the density map (Fig. 2h-n). Subsequent model building identified E8 and E6 as the viral proteins forming the collar and coat-rings, respectively, while L3 constitutes both the skirt and petticoat rings.

### The E6 hexamer forms the coat-ring

Previous studies have implicated E6 as a potential component of the portal complex^2^. Its molecular weight is consistent with that estimated for the coat-ring, and its reported interactions with palisade proteins p4a and A4^11^ align with the observed structural connectivity between the pore and surrounding palisades. To evaluate the structural role of E6, we predicted its monomeric structure by AlphaFold3 (Supplementary Fig. 4a). The prediction revealed two well-defined domains connected by regions of lower confidence. We designated the high-confidence regions as Domain I (residues 172-298) and Domain II (residues 1-161 and 410-567) (Supplementary Fig. 4a,b). Fitting these domains into the STA map demonstrated strong agreement with the upper and lower regions of the coat-ring (Supplementary Fig. 4c,d), accounting for the majority of the observed density (Supplementary Fig. 7a). These findings support the conclusion that E6 constitutes the primary structural framework of the coat-ring.

However, the predicted E6 hexamer exhibited reduced confidence in the upper region (residues 162-409), which adopted an outward-splayed configuration (Supplementary Fig. 4e). Electrostatic analysis indicated that this region is enriched in negatively charged residues (Supplementary Fig. 4f), suggesting that stabilizing interactions with the collar-ring may be required to maintain the compact hexameric conformation observed in our STA map.

### The E8 hexamer constitutes the collar-ring

To identify the proteins forming the collar-ring, we first compiled known E6-interacting proteins and evaluated their interactions with core wall components (Supplementary Table 2). Candidate selection followed three criteria. First, proteins with established structural roles elsewhere in the virion, including p4a, A4, G9^12^, and A27^12,13^, were excluded. Second, volumetric analysis indicated that the collar ring occupies approximately 40% of the coat ring volume, corresponding to an estimated protein length of ∼200-340 amino acids (i.e., ∼35-60% of E6’s length). We used a higher upper threshold to accommodate potential disordered regions. Third, because the collar ring protrudes substantially above the palisade layer and exhibits minimal contact with the inner wall, proteins known to interact strongly with p4b were excluded^1^.

Application of these criteria has shortlisted the candidates to TOP1, MCE, and E8 (Supplementary Table 2). TOP1^14^ and MCE^15,16^ are well-characterized enzymes, whereas E8 lacks a defined function. AlphaFold3 predictions strongly supported E8 as the collar-ring component: the predicted E8 hexamer exhibited markedly superior agreement with the STA density compared with TOP1 or MCE (Supplementary Fig. 5a-d). Moreover, the E8 model displayed uniformly high prediction confidence, consistent with structural stability. Electrostatic surface analysis further revealed a positively charged interface at the base of the E8 hexamer oriented toward the E6 coat-ring (Supplementary Fig. 5e). Collectively, these observations indicate that E8 forms the collar-ring.

### L3 forms both the skirt and petticoat-rings

We next sought to identify the components of the basal region of the pore, comprising an outer skirt-ring and an inner petticoat-ring. In the STA map, the top of the skirt-ring directly interacts with the bottom of the E6 hexamer, while its lower rim contacts the inner wall formed by p4b. Based on these spatial constraints, candidate proteins were restricted to those reported to interact with both E6 and p4b. Volumetric analysis estimated the skirt-ring to occupy approximately 48% of the coat-ring volume. To allow for unresolved or disordered regions, we excluded proteins containing fewer than 200 amino acids. Six proteins met these criteria: ATI (also called A25), I6, M1, L3, ETF1 (also called D6) and DNLI (also called A50) (Supplementary Table 2).

Through systematic prediction of the monomeric and hexameric structures formed by the candidates, L3 emerged as the only model consistent with the experimental structure (Supplementary Fig. 6a). The predicted L3 monomer fits well into each segmented skirt-ring subunit, except for a flexible N-terminal region extending outward (Supplementary Fig. 6b). Interestingly, the predicted L3 hexamer aligned closely with the petticoat-ring but not the skirt-ring density (Supplementary Fig. 6c), suggesting that L3 adopts distinct conformations to form both rings within the pore assembly.

### E8, E6, and L3 assemble into an interwoven pore architecture

Together, these analyses identify E8, E6, and L3 as the principal structural components of the portal pore, forming the collar, coat, skirt, and petticoat-rings. To characterize the molecular organization and channel properties of the pore, we constructed an atomic model of the heteromeric E8-E6-L3 complex (E8:E6:L3 = 6:6:12) (Fig. 2h-j; Supplementary Fig. 7). The model exhibited strong agreement with the STA density map, resolving both secondary structural features and bulky side chains (Fig. 2k-n). Aside from the central plug-like density, minimal unassigned density remained, supporting the assignment of E8, E6, and L3 as the dominant pore constituents.

To visualize inter-subunit interactions, we generated an unfolded schematic representation of the pore (Fig. 3a), in which each ring is depicted as a horizontal strip of six subunits. This representation facilitates side-by-side comparison of individual subunit structures and their lateral interactions. The diagram highlights the staggered arrangement of the 24 subunits across layers, strenghthening inter-subunit packing and structural continuity (Fig. 3).

**Fig. 3.**
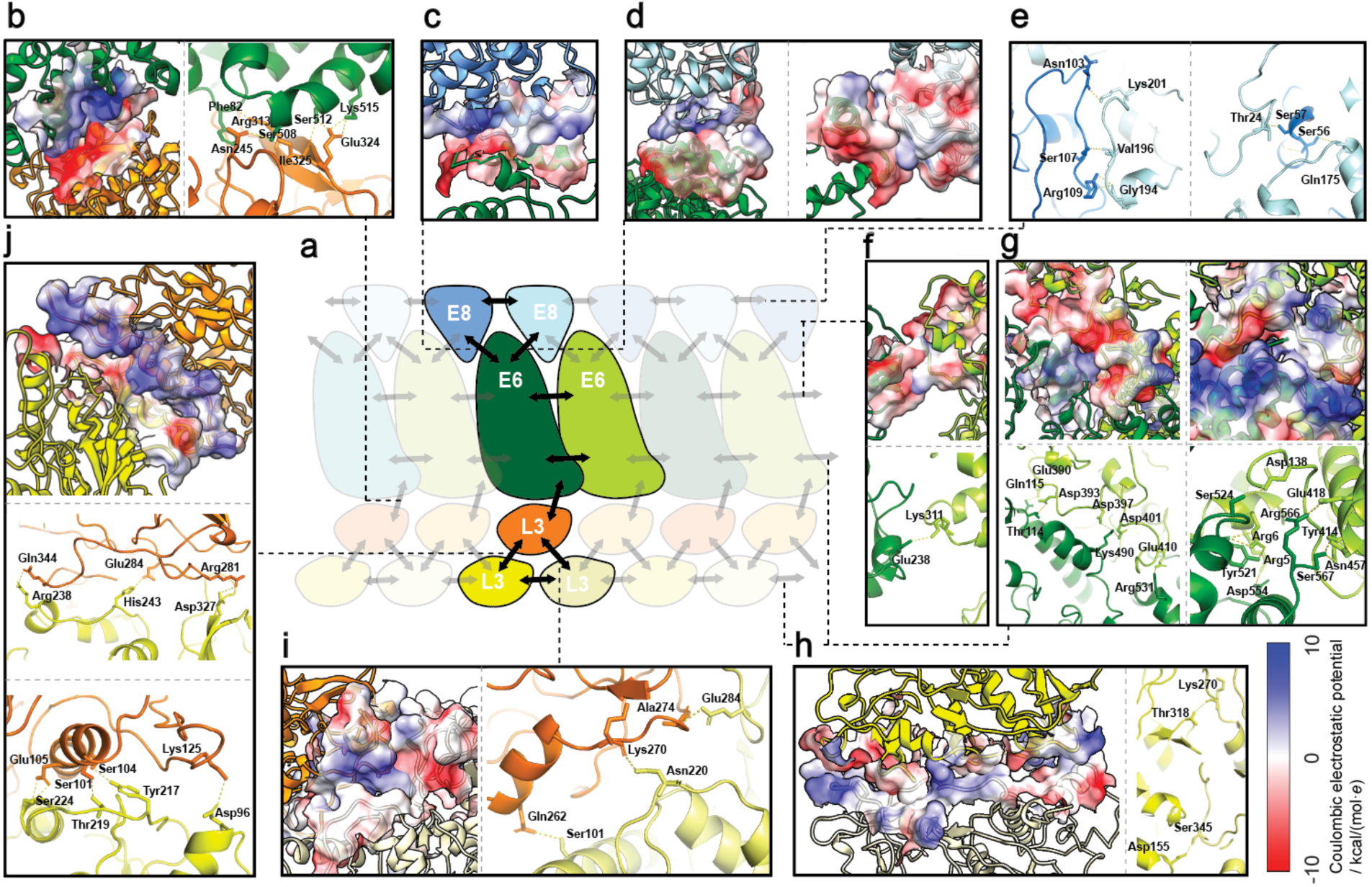
Atomic model reveals the network interaction among pore components. **a**, Unfolded schematic depicting the interaction network among E8, E6, and L3 proteins within the central pore assembly. Each horizontal strip represents a hexameric ring of protein subunits, highlighting lateral interactions and interlayer staggered arrangement for structural stability. **b-j**, Detailed interacting surface between E6 and L3 (**b**); E6 and E8 (**c, d**); adjacent E8 subunits (**e**); upper interfaces between E6 subunits (**f**); lower interfaces between E6 subunits (**g**); and L3 subunits in skirt and petticoat layers (**h-j**). Electrostatic charge distributions are visualized as semi-transparent surfaces, colored according to the charge scale (bottom-right). Hydrogen bonds are represented by yellow bars connecting interacting residues.

The central E6 coat-ring serves as the structural hub of the portal complex. Its upper surface, enriched in negatively charged residues, interacts with adjacent E8 subunits (Fig. 3c,d), while its lower surface engages L3 subunits of the skirt-ring (Fig. 3b). Peripheral interactions between E6 and the palisade protein A10 further anchor the pore within the core wall. Within the basal layer, each L3 subunit of the skirt-ring interacts with two neighboring L3 subunits of the petticoat-ring, stabilizing the lower assembly (Fig. 3i,j). Most interacting surfaces exhibit broadly distributed electrostatic charge complementary to their interacting partners, and possesses multiple interacting pairs potentially forming hydrogen bonds.

Intra-layer interactions further reinforce pore stability. Neighboring E8 subunits engage through multiple potential hydrogen bonds (Fig. 3e). In the E6 coat-ring, each subunit engages its neighbors through a short interface on the upper part and a long interface on the lower part. Both interfaces are covered by complementary charges and potential hydrogen bonds (Fig. 3g).Within the L3 bottom layer, adjacent petticoat subunits also interact through complementarily charged surfaces and potential hydrogen bonds (Fig. 3h), while no direct contact is observed between neighboring skirt-ring subunits (Fig. 2f).

Overall, the pore assembly is stabilized predominantly by extensive electrostatic complementarity and hydrogen bonding. The robust inter-subunit interactions within the L3 basal layer (Fig. 3h-j) correlate with the highest local resolution observed in this region (Supplementary Fig. 3b), whereas weaker interactions in the upper E6 region (Fig. 3f) are consistent with reduced local resolution, suggesting increased conformational flexibility.

### The pore contributes to early transcription within the viral core

Although the molecular mechanisms remain incompletely understood, all three pore components identified here have previously been implicated in the formation of transcriptionally competent cores. Early genetic studies of conditional lethal mutants (Dts^17^ and Cts^18^ collections) demonstrated that strains harboring mutations in the E8R gene^19^ (Dts23 and Cts19) and the E6R gene^20^ (Cts52) produce progeny virions that are morphologically indistinguishable from wild-type particles and retain host entry capability, yet exhibit profound defects in early transcription¹⁷˒¹⁸. Similarly, conditional suppression of L3 expression results in virions that preserve structural integrity but are impaired in early gene transcription^9^.

These observations collectively support a model in which E8, E6, and L3 are dispensable for virion morphogenesis but essential for early transcriptional activity. Notably, none of these proteins are components of the established transcription machinery responsible for mRNA biogenesis^21^, suggesting that their functional contribution occurs at a mechanistically distinct stage. Moreover, despite the transcriptional deficiencies observed in intact mutant cores, viral extracts derived from E8R temperature-sensitive mutants grown under non-permissive conditions and virions lacking L3 remain transcriptionally active *in vitro*^9,19^. This discrepancy indicates that the functional requirement for E8, E6, and L3 is specific to the structural context of intact cores, rather than intrinsic catalytic processes of transcription.

Our structural analysis reveals that the pore constitutes the largest continuous channel observed within the core wall, directly connecting the interior of the viral core with the host cytoplasm. When considered alongside prior functional studies, these findings suggest that the pore mediates the exchange of molecules required for early transcription.

### A lysine-lined tube within the L3 layer forms the central constriction of the pore

To quantitatively evaluate the transport capacity of the pore, we analyzed the geometry of the central channel using CAVER 3.0 (Fig. 4a,b). The channel extends more than 18 nm along the longitudinal axis of the pore, traversing the L3, E6, and E8 layers (Fig. 4a). Within the L3 region, three discrete constriction sites define the narrowest segment of the channel (Fig. 4b-e). These constriction sites contain a series of lysine residues (Lys261, Lys268, Lys270, and Lys272), which project inward to create a ∼2-nm-long, positively charged tube (Fig. 4b). The minimal diameters of the constriction sites measure approximately 17-19 Å. These dimensions are sufficient to accommodate single-stranded RNA and small metabolites, yet are incompatible with the passage of double-stranded DNA (Fig. 4b). Given that most cytosolic DNA sensors, such as cGAS, IFI16, and AIM2, possess a net positive charge and have a diameter exceeding 1.7 nm, their passage through the channel is highly improbable. In summary, the transport capacity of this lysine-lined tube aligns with the expected permeability of the core wall.

**Fig. 4.**
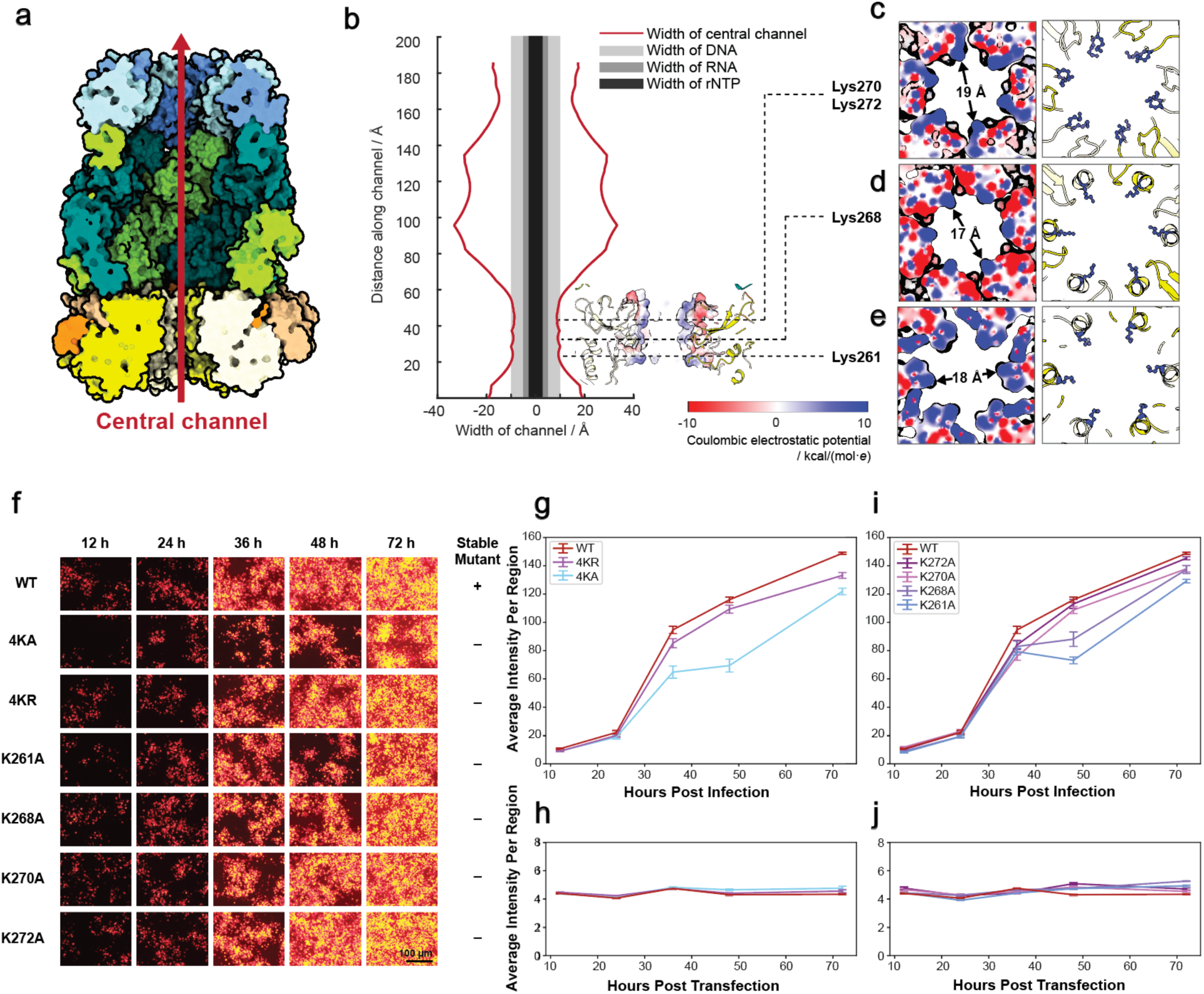
The lysine-lined tube within the pore and its functional importance. **a**, Cross section showing the central channel within the E8-E6-L3 complex. **b**, Channel width profile calculated using CAVER 3.0, plotted against distance along the channel. Red line represents the central channel, Light grey, grey, and dark grey bars indicate widths of DNA, RNA, and ribonucleotide molecules, respectively. Dashed lines highlight constriction sites within the central channel. **c-e**, Enlarged top view of the three constriction sites marked by dashed lines in panel **b**. Diameters at the narrowest points are indicated by black arrowheads. Constriction residues are highlighted in blue. Electrostatic charge distribution on the constriction sites is shown in color blue and red according to the color key in panel **b**. **f**, Fluorescence images of cells during mutant virus rescue experiments. Fluorescent images were recorded at 12, 24, 36, 48 and 72 hours after plasmid transfection and template virus infection. **g**, Fluorescence intensity measured during WT, 4KR and 4KA mutant virus rescue. **h**, Fluorescence intensity measured during WT, 4KR and 4KA plasmid transfection. **i**, Fluorescence intensity measured during WT, K272A, K270A, K268A and K261A mutant virus rescue. **j**, Fluorescence intensity measured during WT, K272A, K270A, K268A and K261A plasmid transfection. The scalebar in **g-j** represents the standard error of mean (sem).

Sequence conservation analysis further indicates functional importance of this lysine-lined tube. Across the *Orthopoxvirus* genus, the L3 protein exhibits high sequence conservation, with an average identity of 93% (Supplementary Fig. 8a). Even among more distantly related members of the *Chordopoxvirinae*, residues corresponding to VACV Lys261, Lys268, Lys270, and Lys272 overwhelmingly retain positive charges. Notably, all viral sequences examined at UniProt^22^ contained at least three positively charged residues at these positions.

Despite substantial sequence divergence, AlphaFold3 predictions indicate that L3 homologs from diverse genera adopt structurally similar hexameric assemblies resembling the petticoat-ring observed in VACV (Supplementary Fig. 8b-e). In each case, residues corresponding to the VACV constriction lysines line a positively charged channel of comparable dimensions. This combined structural and electrostatic conservation supports a conserved functional role for the lysine-lined tube.

### Functional importance of the lysine-lined tube in virus replication

If the E8-E6-L3 pore functions as a selective conduit for RNA export, residues lining the channel constriction would be expected to play critical roles in viral gene expression. To test this hypothesis, we generated recombinant viruses in which the four lysine residues (K261, K268, K270, and K272) were substituted with alanine (4KA) or arginine (4KR). Substitution with alanine eliminates positive charges and alters the chemical environment of the channel interior, whereas arginine substitutions preserve positive charges while slightly narrowing the pore diameter. To monitor viral gene expression, an mCherry reporter gene was inserted downstream of the mutated L3 locus.

Consistent with expectations, all virus rescue groups exhibited fluorescence signals markedly exceeding those of plasmid transfection-only controls (Fig. 4f-j; Supplementary Fig. 9). However, beginning at 36 hours post-infection, fluorescence accumulation in the 4KA mutant was significantly reduced relative to the wild-type (WT) control, which contained only the reporter gene insertion. In comparison, the 4KR mutant exhibited only a modest delay (Fig. 4f, g). These findings indicate that electrostatic properties of the constriction region exert a greater functional influence than channel diameter alone.

To further dissect residue-specific contributions, we generated viruses carrying individual alanine substitutions at each lysine position (K261, K268, K270, or K272). Among these, the K261A mutant exhibited the most pronounced reduction in fluorescence intensity after 48 hours post-infection (Fig. 4i), suggesting a critical role for this residue. Notably, except for the WT control group, none of the mutant viruses yielded stable progeny upon amplification despite successful initial rescue. This observation indicates that all four lysine residues are essential for productive VACV replication.

Collectively, these results infer a critical role for the lysine-lined tube in the replication of VACV and other poxviruses, likely through its involvement in the material exchange during early transcription.

## Discussion

Early transcription within the poxvirus core represents a defining and mechanistically distinctive feature of poxvirus biology. Although recent studies have significantly advanced our understanding of viral core organization^1–3^ and the architecture of the transcription machinery^15,23–25^, the structural basis underlying molecular exchange across the core wall has remained unresolved. The present study addresses this gap by resolving the structure of the VACV portal complex and establishing a near-complete atomic model of its pore.

Our analysis reveals that the pore is assembled from three viral proteins, E8, E6, and L3, organized into a multi-layered, hexamerically symmetric architecture embedded within the core wall. Importantly, the structure was determined from intact, envelope-free viral cores that closely resemble the native post-entry state, thereby providing a structural framework for investigating transcriptionally competent cores. This apo-state model offers a foundation for future studies aimed at elucidating dynamic conformational changes associated with active transcription.

The overall architecture of the VACV portal complex closely resembles the lower-resolution MPXV portal structure we previously reported^6^. Given the substantial genomic and proteomic conservation among *Orthopoxvirus* genus, the structural principles described here are likely applicable to other pathogenic members, including VARV and MPXV.

The central channel’s constriction sites within the VACV pore are markedly enriched with positively charged lysine residues. This region is not only functionally critical for VACV replication but also exhibit striking structural and electrostatic conservation across genera, presenting these sites as potential new targets for antiviral intervention. Notably, both the geometry and electrostatic characteristics of the constriction region resemble those of the SARS-CoV-2 nsp3-nsp4 pore complex^26^, which mediates RNA transport across double-membrane vesicles. This similarity suggests a potentially analogous function govern material translocation in structurally distinct viral systems.

In the VACV pore, while the constriction region is defined by L3, the E6 and E8 layers form expanded chambers containing a plug-like density of currently unknown identity. The presence of this density raises the possibility that the portal complex may incorporate regulatory or gating mechanisms beyond passive transport. Although our findings strongly support a functional role for the pore in molecular exchange, the precise mechanistic details of substrate recognition, transport dynamics, and gating remain to be determined. In particular, identification of the plug-like density and characterization of potential conformational rearrangements during active transcription represent important directions for future investigation.

Lastly, our work highlights the power of cryo-ET and STA in resolving high-resolution complex structures from thick biological specimens (>300 nm)^27,28^. The methodological advances demonstrated here provide a strategic framework for investigating macromolecular assemblies that remain inaccessible to conventional single-particle cryo-EM approaches, particularly within complex viral systems.

## Materials and Methods

### VACV infected cell sample preparation

For the *in-situ* sample preparation, Vero cells were cultured on 200 mesh continuous carbon film coated gold grids (EMCN, CN) over night. After the cell is fully attached to the grid, we transferred the grid to a new dish to incubate with VACV at multiplicity of infection (MOI) at 25 in Dulbecco’s modified eagle medium (DMEM) for 1 hour at 37 ℃. Then the grid is transferred to DMEM supplemented with 10% fetal bovine serum (FBS) and 1% antibiotics (10,000 IU/ml penicillin and 10,000 µg/ml streptomycin), and cultured at 37 ℃ in 5% CO_2_ for 24 hours. For cryo-ET sample preparation, the grid was supplemented with 4 µl DMEM and blotted for 12 seconds before plunge frozen in liquid ethane using a Cryo-plunger 3 (Gatan, CA). Infectious sample was handled in the BSL-2 Facility, School of Life Sciences, Tsinghua University.

### VACV core sample preparation

For virus propagation, Vero cells were cultured in six T-175 flasks with DMEM supplemented with 10% FBS and 1% antibiotics (10,000 IU/ml penicillin and 10,000 µg/ml streptomycin) at 37 ℃ in 5% CO_2_. Cultured Vero cells were infected with a MOI at 0.01 by VACV strain Western Reserve (WR) after the cell reaches 90% confluence. The infected cells were then cultured in DMEM supplemented with 2.5% FBS and 1% antibiotics for 3 days at 37 ℃ in 5% CO_2_ before harvest.

On three-day post-infection, all cells exhibited significant cytopathic effect while most cells remained attached to the flask bottom. The infected cells were detached from the flask bottom and isolated. Then the cell pellets were resuspended in phosphate-buffered saline (PBS) and lysed with a no-touch ultrasonic homogenizer (SCIENTZ, CN) at 80% power. The MVs were pelleted from supernatant by ultracentrifugation (Beckman, IN). Then, the pellet containing MVs were treated with NP40 and dithiothreitol (DTT) to dissolve the viral envelope. Then, released viral cores were isolated and purified through sucrose cushion and sucrose-density gradient by ultracentrifugation (Beckman, IN). The fractions-to-collect were changed into sucrose-free buffer for cryogenic sample preparation. VACV core samples were supplemented with BSA-coated gold tracer (Aurion, NED) before plunge freezing. All infectious samples were handled in the BSL-2 Facility, School of Life Sciences, Tsinghua University.

### Cryo-electron tomography and data preprocessing

The cryogenic sample was imaged using a 300 kV Titan Krios electron microscope (Thermo Fisher Scientific, Hillsboro, OR) equipped with GIF Quantum energy filter (slit width 20 eV) and K3 direct electron detector (Gatan, CA). All tilt series were collected using dose-symmetric scheme in SerialEM^29^ with settings in Supplementary Table 1.

Preprocessing of the raw data is automatically done using an in-house developed software^30^. In brief, the first frame in each movie was removed and the last 9 frames were motion corrected and averaged by MotionCor2^31^. Then, defoci of the averaged movies were estimated using Gctf^32^ and the tilt series were aligned using AreTomo 1.2.5^33^ or ETomo^34^. The tilt images were then dose-weighted and three-dimensional contrast transfer function (CTF) corrected and reconstructed into tomograms with NovaCTF^35^. For visualization purposes, tomogram slices of purified core were denoised using cryo-CARE^36^, and tomogram slices of *in-situ* cores were low-passed to 80 Å.

### Sub-tomogram averaging

The tomograms were 8-fold binned for particle picking with PyTom^37^ using EMD-18917 as template. Sub-tomogram averaging was done in RELION-4.0^38^. Initially, 22,808 particles were identified from 248 tomograms by template matching, and subsequently classified at 8-fold binning. 3,035 particles were removed as misidentified particles during this step, and the remaining particles were classified into two groups: one group containing 4,810 particles resulted in an optimal reconstruction, and the other group containing 14,963 particles resulted in a suboptimal reconstruction. The optimal 4,810 particles were refined globally to a map at Nyquist resolution (10.88 Å) at 4-fold binning, which was subsequently used as a template for further classification of the suboptimal group. 3,169 particles were found from the suboptimal group and merged with the initial optimal 4,810 particles for further refinement at 2-fold binning. To obtain the structure of the intact portal complex, the 7,979 particles were refined with a large mask until the refinement converged at 7.1 Å resolution. The final portal complex structure was reconstructed from 2D images at 2-fold binning and sharpened using a B-factor of -294.205 Å^2^. To achieve higher resolution of the pore, we masked away the surrounding palisades and the central plug densities for further refinement at 2-fold binning, followed by frame alignment at 1-fold binning. The final structure achieved 4.9 Å resolution, was reconstructed from 2D images and sharpened using a B-factor of -122.349 Å^2^.

### Density map segmentation and building the composite structure of the viral core

To build an intact virus core model, the core wall and interior density were segmented manually from cryo-CARE^36^ processed tomograms using 3D Slicer^39^ at 8-fold binning. Palisade proteins were oversampled and STA aligned using Dynamo^40^ at 8-fold binning to Nyquist resolution. Duplicated particles and misaligned particles were removed according to particle distance, cross correlation and geometric restraints. The portal complexes and palisades were projected to their refined coordinates relative to the segmented core wall and interior densities. Image and video rendering of the model is done in ChimeraX^41^.

### Statistical analysis

For the statistical analysis of portal complex particle number, 46 virus particles intactly included in the tomograms were randomly selected. Then, we manually labeled the boundary of each virus core and each particle with final refined coordinates with in this boundary was counted for one portal complex on this virus core. For pore orientation analysis, we manually defined the short axis of the same 46 virus cores in IMOD^42^ as the vector which is vertical to the largest section of the virus core and passes the center of the largest section. Orientation of each portal complex was defined as the orientation of the axis of C6 symmetry. Each particle’s relative orientation to short axis is calculated and plotted. A total of 904 portal complexes were included in the statistical analysis above.

### Atomic model building for the pore

AlphaFold3^43^ -predicted models of the E6 domains, full-length E8 hexamer and structured region of the L3 hexamer fit well into the STA density maps of the coat-ring (Supplementary Fig. 7a), collar-ring (Supplementary Fig. 7b) and petticoat-ring (Supplementary Fig. 7c). However, it failed to predict the correct hexameric conformations of full length E6 and L3 in the skirt-ring conformation.

To obtain an initial model of the upper half of the pore, we predicted a heterododecamer comprising six E6 fragments (residues 162-409) and six full-length E8 subunits. The resulting model aligned well with the STA density (Supplementary Fig. 7d). Similarly, to obtain an initial model of the lower half of the pore, we predicted a second heterododecamer comprising six E6 domain II (residues 1-161 and 410-567) and six L3 structured fragments (residues 62-350). In this context, L3 was predicted to adopt a conformation that fit well within the skirt-ring density, with only minor discrepancies (Supplementary Fig. 7e).

The E6-E8 and E6-L3 models were then flexibly fitted to their corresponding densities using MDFF^44^ in VMD^45^ (Supplementary Fig.7d-e). Combining the two flexibly fitted models and the predicted L3 petticoat hexamer, we obtained an initial composite model of the pore. While the global fit was satisfactory, this model remained fragmented and exhibited local discrepancies (Supplementary Fig. 7f). To resolve these, we manually connected the domains and refined the model using Coot^46^ and PHENIX^47^, correcting for local inaccuracies (Supplementary Fig. 7g, Supplementary Table 3).

In summary, the atomic model was built in three steps. First, we fragmented the pore into three parts for Alphafold3^43^ prediction. The upper part includes 6 copies of full length E8 and 6 copies of E6 (residues 162-409). The middle part includes 6 copies of E6 (residues 1-161), 6 copies of E6 (residues 410-567), and 6 copies of L3 (residues 62-350). The lower part includes 6 copies of L3 (residues 62-350). Second, the three predicted models were flexibly fitted to their corresponding densities segmented from the pore structure using MDFF^44^ in VMD^45^ and combined together as an initial model. Third, the Initial model was manually connected using Coot^46^, and refined using PHENIX real-space refinement^47^.

### Cavity and sequence analysis

Cavity analysis of the pore is performed using CAVER 3.0^48^. The output result from CAVER 3.0 was imported to Matlab for further analysis and visualization. Sequence alignment across different species and genus is done using UniProt^22^ blast and align tools.

### Construction of recombinant plasmids

To generate recombinant vaccinia viruses, transfer vectors for homologous recombination were constructed. Each vector contained an mCherry fluorescent reporter gene under the control of a vaccinia early/late (E/L) promoter, which served as a selection marker inserted between the G9R and L1R genes, downstream of the reverse-oriented L3L gene. Homology arms targeting the viral genome, which consisting of a 663 bp left arm (partial L3L sequence) and a 666 bp right arm (partial G9R sequence), were designed to facilitate targeted insertion. The recombination cassettes were cloned into a pUC57 backbone, yielding six recombinant plasmids harboring mutations K261A, K268A, K270A, K272A, 4KA, and 4KR, along with an additional control plasmid. All constructs were verified by restriction digestion and sequencing prior to transfection.

### Virus rescue, fluorescence imaging and image analysis

Mutated virus rescue was applied with vaccinia virus strain Tiantan (VTT). BHK-21 cells were seeded at 2×10⁵ cells/wall in twelve-well plates. Cell culture medium is replaced by DMEM supplemented with 2% FBS after the cells grow to 80% confluency. For virus rescue experiments, cells were infected with VTT at MOI=0.05 and transfected with donor plasmid carrying mutated gene and reporter gene (1 µg plasmid / well) using X-tremeGENE HP DNA transfection reagent. Each mutant rescue experiment is repeated three times. At 12 h, 24 h, 36 h, 48 h, 72 h post-infection, five fluorescence images were randomly recorded from each group with 100 ms exposure time using Olympus microscope IX73. For the plasmid transfection-only control group, donor plasmid was transfected and imaged parallelly to the virus rescue groups, but not infected with VTT.

Using ImageJ^49^, we cropped fluorescent images into regions of 800 pixels × 600 pixels and measured the average intensity in each region. The average fluorescent intensity and the standard error of mean (sem) of each group was calculated and plotted using Python.

## Acknowledgements

We thank Dr. Jianlin Lei, Dr. Fan Yang, Dr. Xiaomin Li and Dr. Tao Liu from the cryo-EM Facility, Technology Center for Protein Sciences, Tsinghua University, for their support on cryo-EM data collection. We thank the computational facility support on the cluster of Bio-Computing Platform (Tsinghua University Branch of China National Center for Protein Sciences Beijing). We thank the BSL-2 Facility, Tsinghua University, for their support on BSL-2 sample preparation. We are in debt to Dr. Qiangfeng Zhang from Tsinghua University for advises on model building.

## Declarations

### Author contributions

S.L. conceived and supervised the project. Y.H., K.W., and J.Z. prepared the samples. Y.H., K.W., and Z.Zhang collected and processed the EM data. Y.H., K.W., C.P. and W.Z. processed the EM data. Y.H., R.L., Z.Zhu, C.S. and S.L. analyzed the structures. S.Liu, Y.H., J.Z. and B.H. performed the functional experiments. Y.H., W.T. and S.L. wrote the manuscript. All authors critically revised the manuscript.

### Date and materials availability

Electron microscopy maps and atomic models of the portal complex and pore have been deposited in the Electron Microscopy Data Bank and Protein Data Bank under accession codes EMD-65763, EMD-65762, and PDB-9W91.

## Supplementary Information

**Supplementary Fig. 1.**
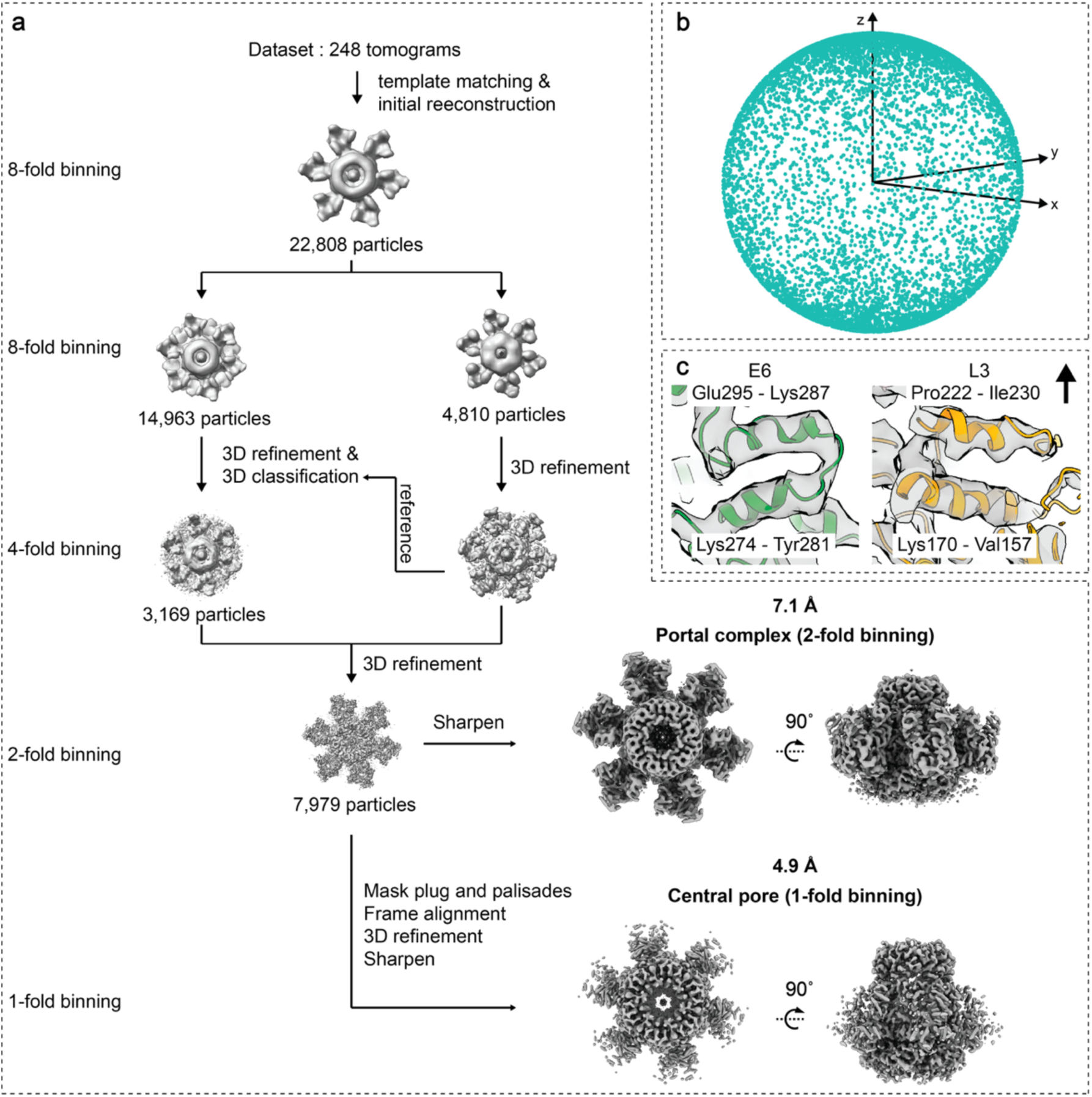
VACV pore STA workflow. **a**, 22,808 particles were identified by template matching from 248 tomograms for STA as shown in the workflow. 7,979 particles were finally classified as portal complex and reconstructed into the structure displayed in main text figure. Structure of the pore was further refined with a narrower mask encompassing the pore exclusively. **b**, Angular distribution of the particles used for final structure reconstruction shows no obvious preferred orientation. **c**, Local zoom-in view of the density map with our atomic model fitted shows no obvious preferred orientation induced “smear effect” along the symmetrical axis. The black arrowhead shows direction of the symmetrical axis.

**Supplementary Fig. 2.**
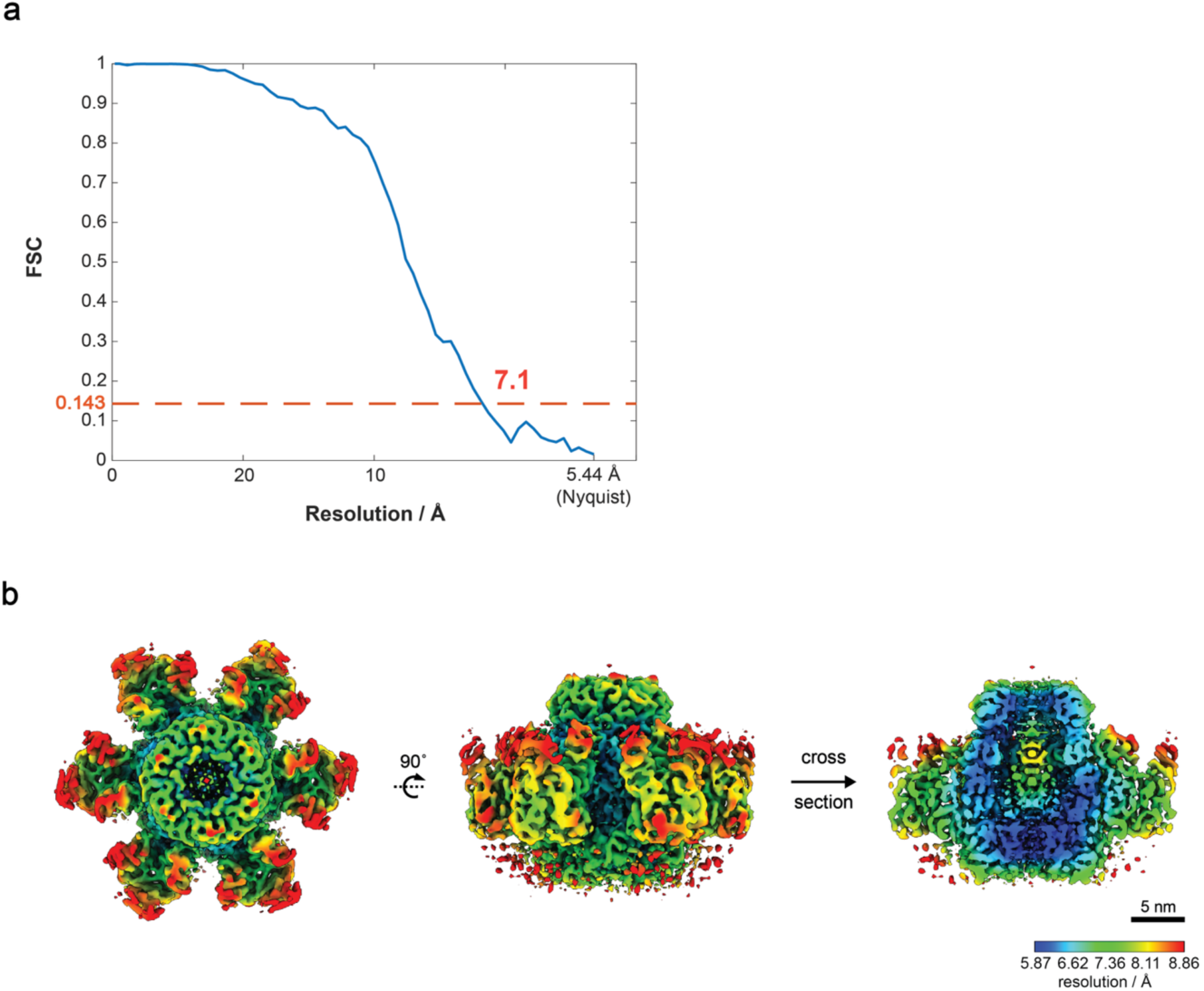
FSC curve and local resolution of portal complex STA structure. **a**, Fourier shell correlation (FSC) curve of the STA structure of the portal complex. **b**, Top, side and cut-open view of the STA structure of the portal complex colored according to local resolution.

**Supplementary Fig. 3.**
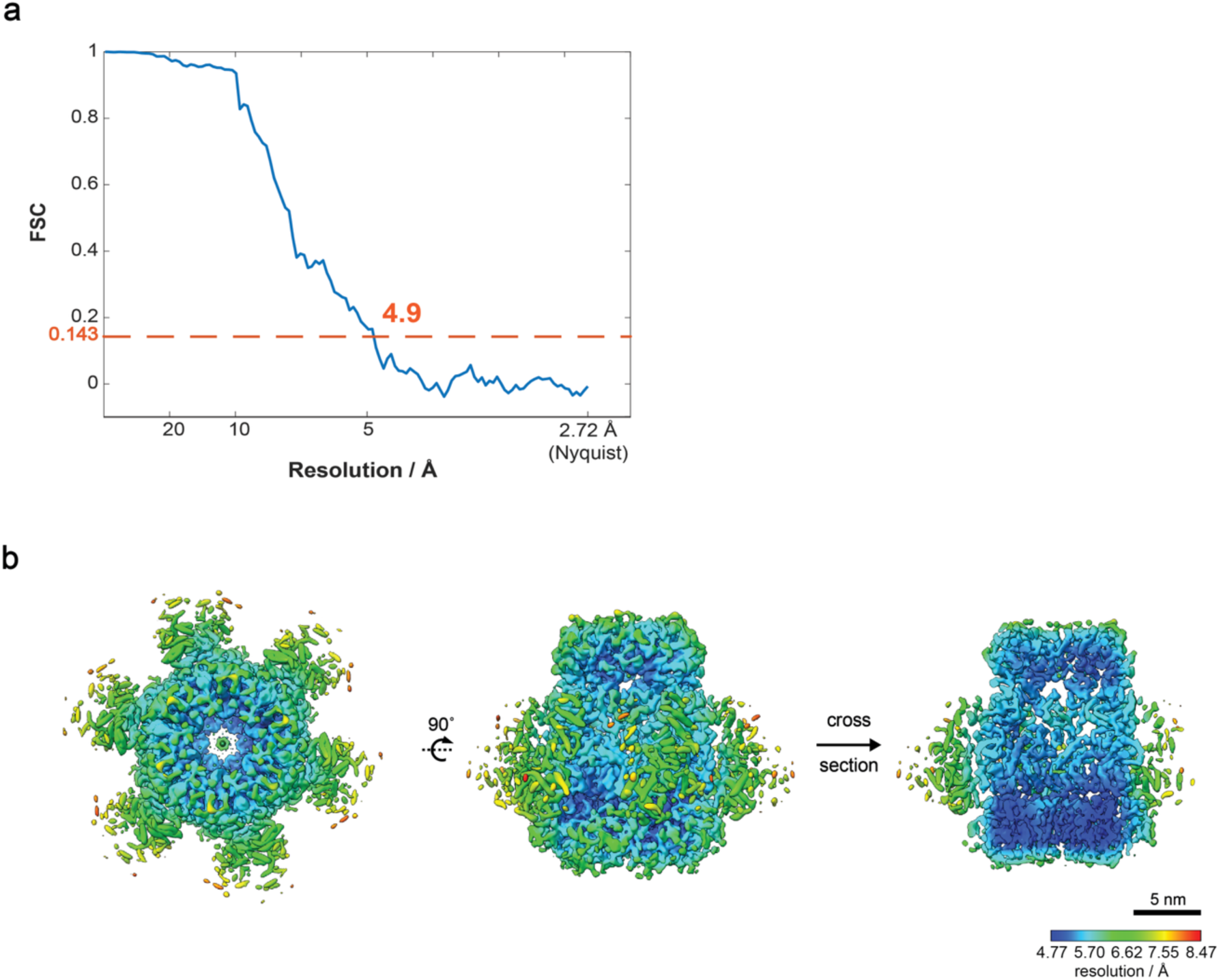
FSC curve and local resolution of pore STA structure. **a**, Fourier shell correlation (FSC) curve of the STA structure of the pore. **b**, Top, side and cut-open view of the STA structure of the pore colored according to local resolution.

**Supplementary Fig. 4.**
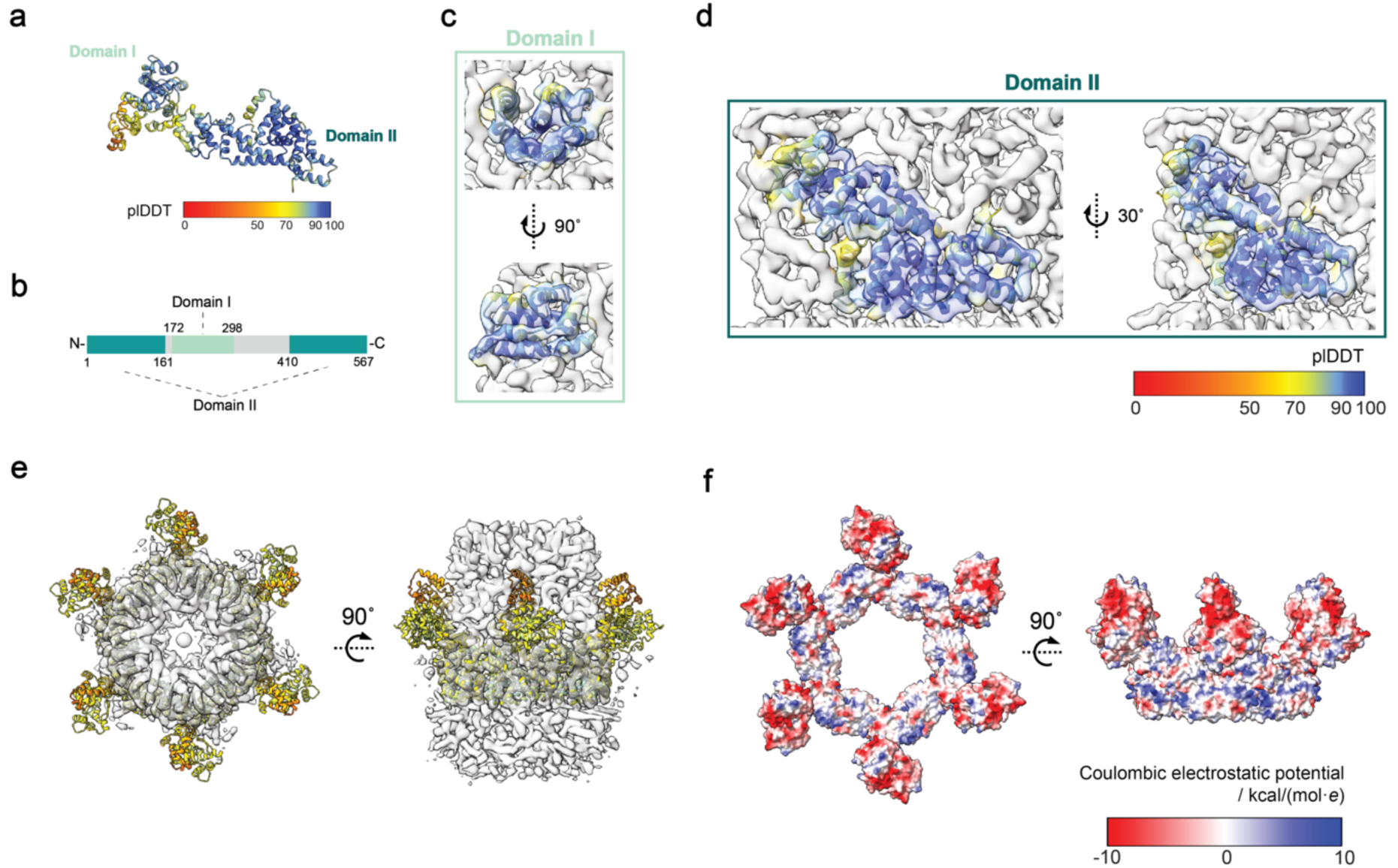
Alphafold3 predicted E6 structure fitted to the pore STA map. **a**, An Alphafold3 predicted E6 monomer. The color corresponds to the plDDT score of the prediction as shown in the color key. **b**, Domian definition of protein E6. Light green marks the amino acid region of domain I and dark green marks the amino acid region of domain II. **c**, The zoom in view of domain I fitted to the pore structure. **d**, The zoom in view of domain II fitted to the pore structure. **e**, The Alphafold3 predicted E6 hexamer is colored according to the plDDT score of the prediction according to the color key below **d** and fitted to the middle ring of the pore. The model is displayed in top view and side view. **f**, Charge distribution of the Alphafold3 predicted E6 hexamer is displayed on the model according to the color key on right lower corner. Red color stands for negative charges and blue color stands for positive charges.

**Supplementary Fig. 5.**
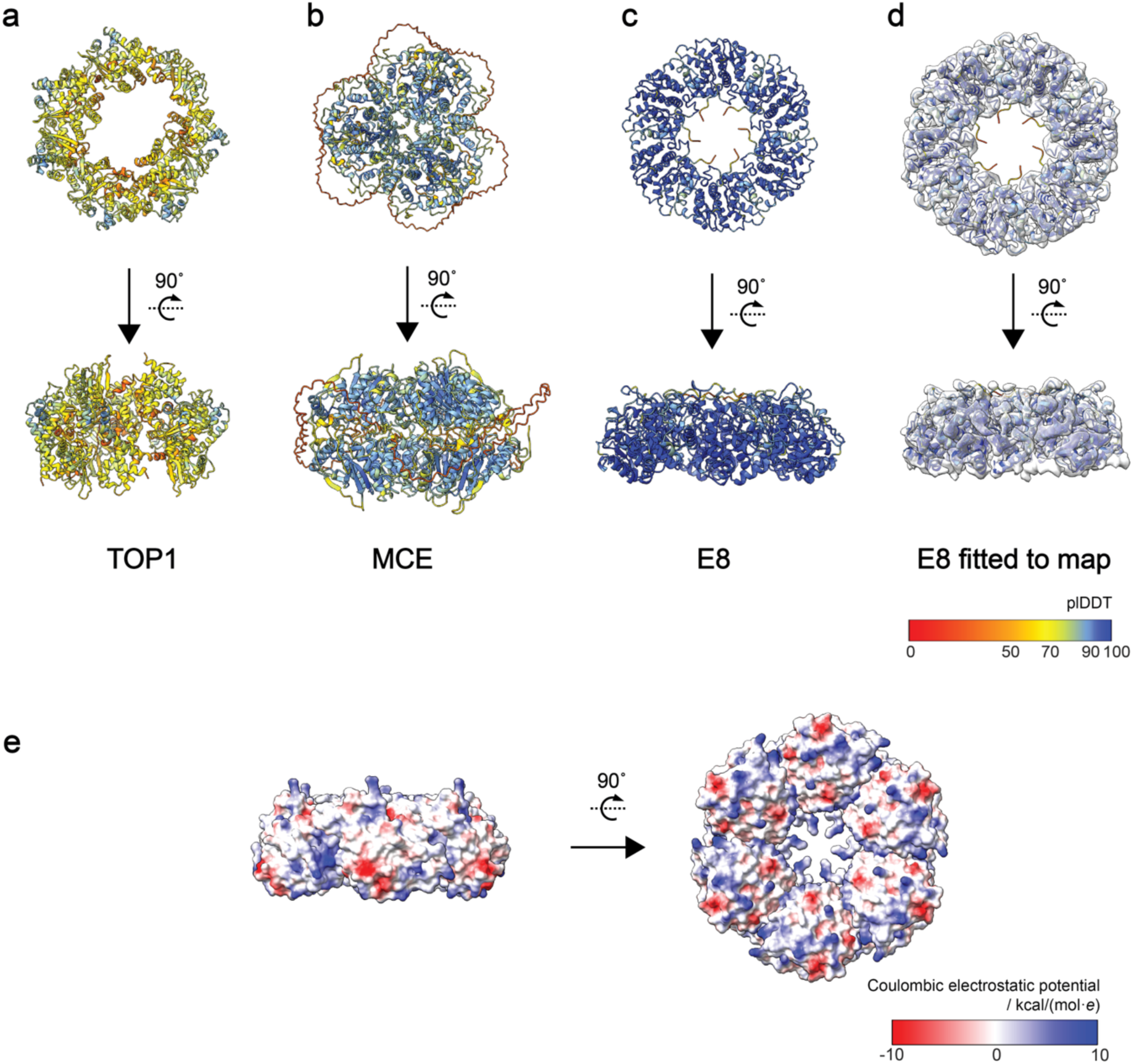
Alphafold3 predicted top ring candidates. **a-c**, AlphaFold3 predicted TOP1, MCE and E8 hexamers are shown in top view and side view. The model is colored according to the plDDT score of the prediction. **d**, AlphaFold3predicted E8 hexamer fitted to the segmented top layer density is shown in top view and side view. **e**, Charge distribution of the Alphafold3 predicted E8 hexamer is displayed on the model according to the color key on right lower corner. Red color stands for negative charges and blue color stands for positive charges. The model is shown from side and bottom.

**Supplementary Fig. 6.**
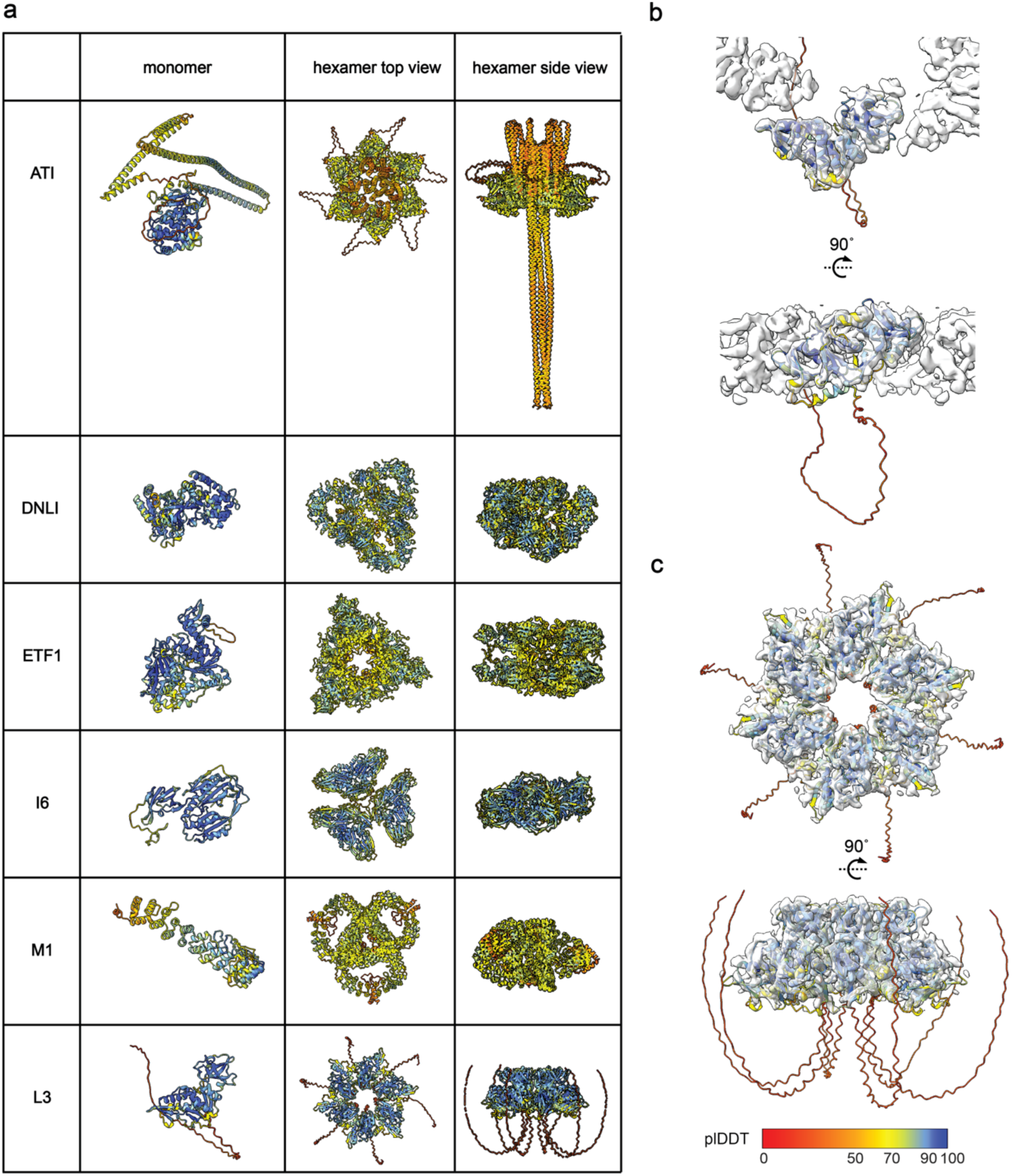
Alphafold3 predicted bottom ring candidates. **a**, Alphafold3 predicted ATI, DNLI, ETF1, I6 and M1monomer and hexamers **b**, Predicted L3 monomer fitted to the segmented skirt-ring. **c**, Predicted L3 hexamer fitted to the segmented bottom layer “petticoat”. The model is colored according to the plDDT score of the prediction.

**Supplementary Fig. 7.**
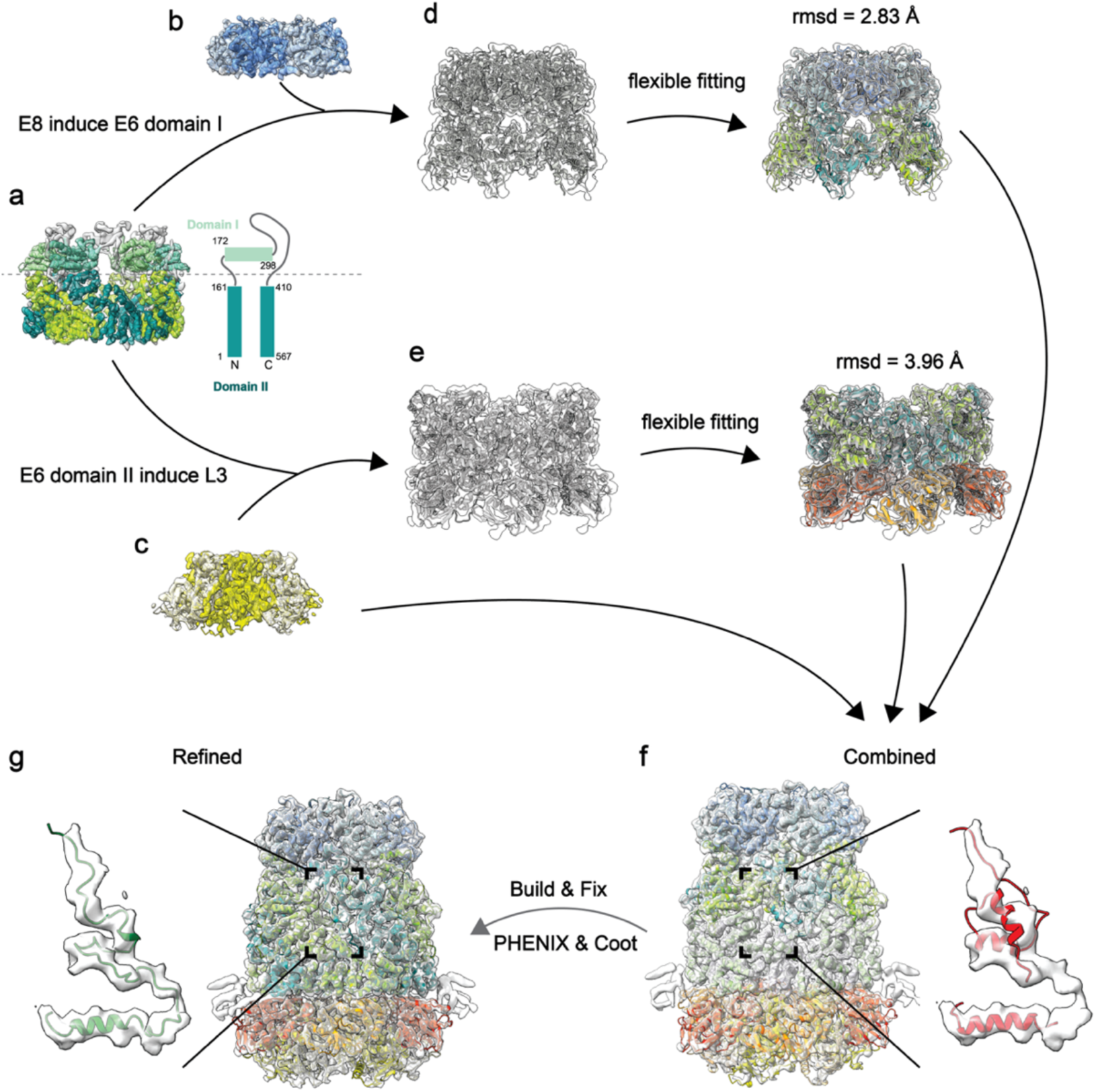
Workflow of pore protein components identification and model building. **a**, Alphafold3 predicted E6 domain I and domain II fitted to the middle layer density of the pore. **b**, Alphafold3 predicted E8 hexamer fitted to the top layer density of the pore. **c**, Alphafold3 predicted L3 hexamer fitted to the bottom layer petticoat density of the pore. The disordered regions were removed. **d**, E6 residue 162-409 predicted with E8 hexamer matches to the top layer and the upper part of the middle layer density. The model fits to density map satisfyingly after flexible fitting with final rmsd = 2.83 Å. **e**, L3 ordered region (residues 62-350) predicted with E6 lower part (residues 1-161 and 410-567) matches to the bottom layer skirt and the lower middle layer density. The model fits to density map satisfyingly after flexible fitting with final rmsd = 3.96 Å. **f**, The combined model of **c**, **d** and **e** flexibly fitted to their corresponding densities. **g**, Final atomic model built and fixed according to the density map. The zoom-in views in f and g shows the same region before and after fixation in PHENIX and Coot.

**Supplementary Fig. 8.**
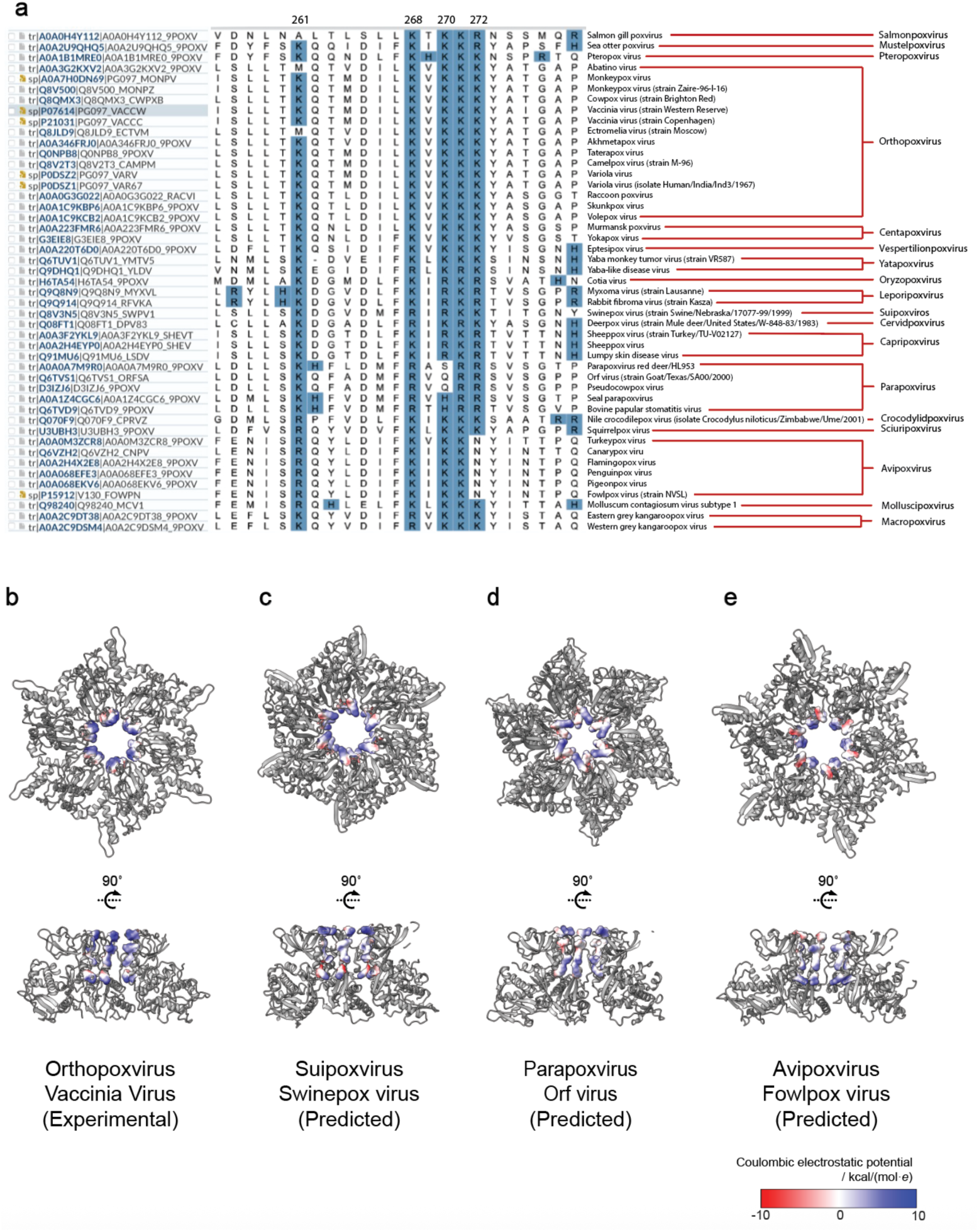
Sequence and predicted structure of L3 homologous protein. **a**, Aligned sequence of L3 homologous protein in representative species selected from *chordopoxvirinae*. Positively charged residues are highlighted in blue. Numbers above label the four constriction residues according to their position in vaccinia L3. **b**, VACV L3 pore petticoat structure shown in top view and cut-open sideview. **c-e**, Structure of hexameric L3 homologous protein in Swinepox virus, Orf virus and Fowlpox virus predicted using Alphafold3 are shown in top view and cut-open sideview. Electrostatic charges on the constriction sites are shown on the half transparent surface according to the color bar on the lower right corner.

**Supplementary Fig. 9.**
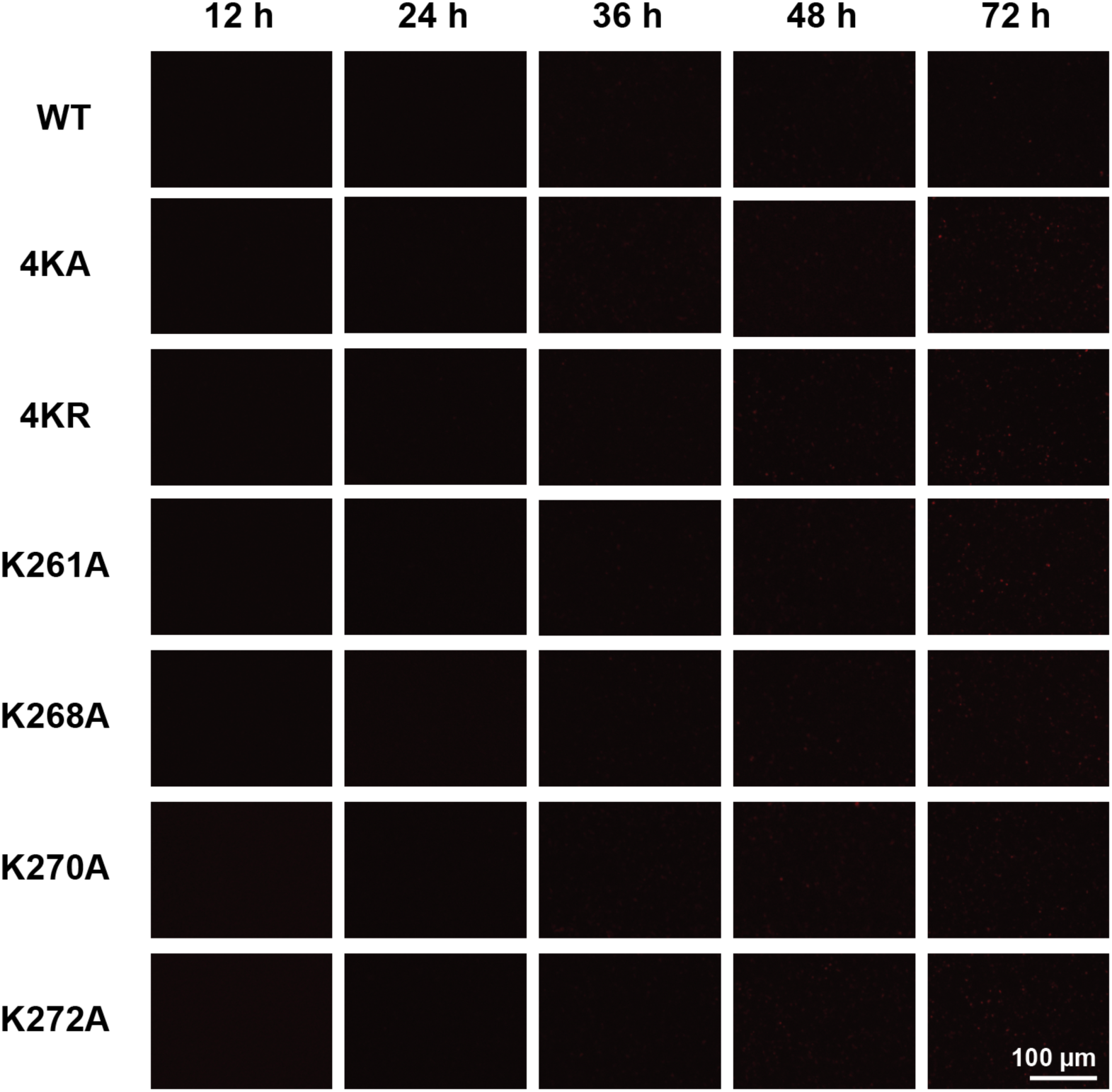
Fluorescence images of the plasmid transfection-only control. The fluorescent images of BHK-21 cells transfected with donor plasmid carrying mCherry reporter gene and mutated L3 gene are imaged at 12 h, 24 h, 36 h, 48 h and 72 h post transfection.

**Supplementary Table 1.**
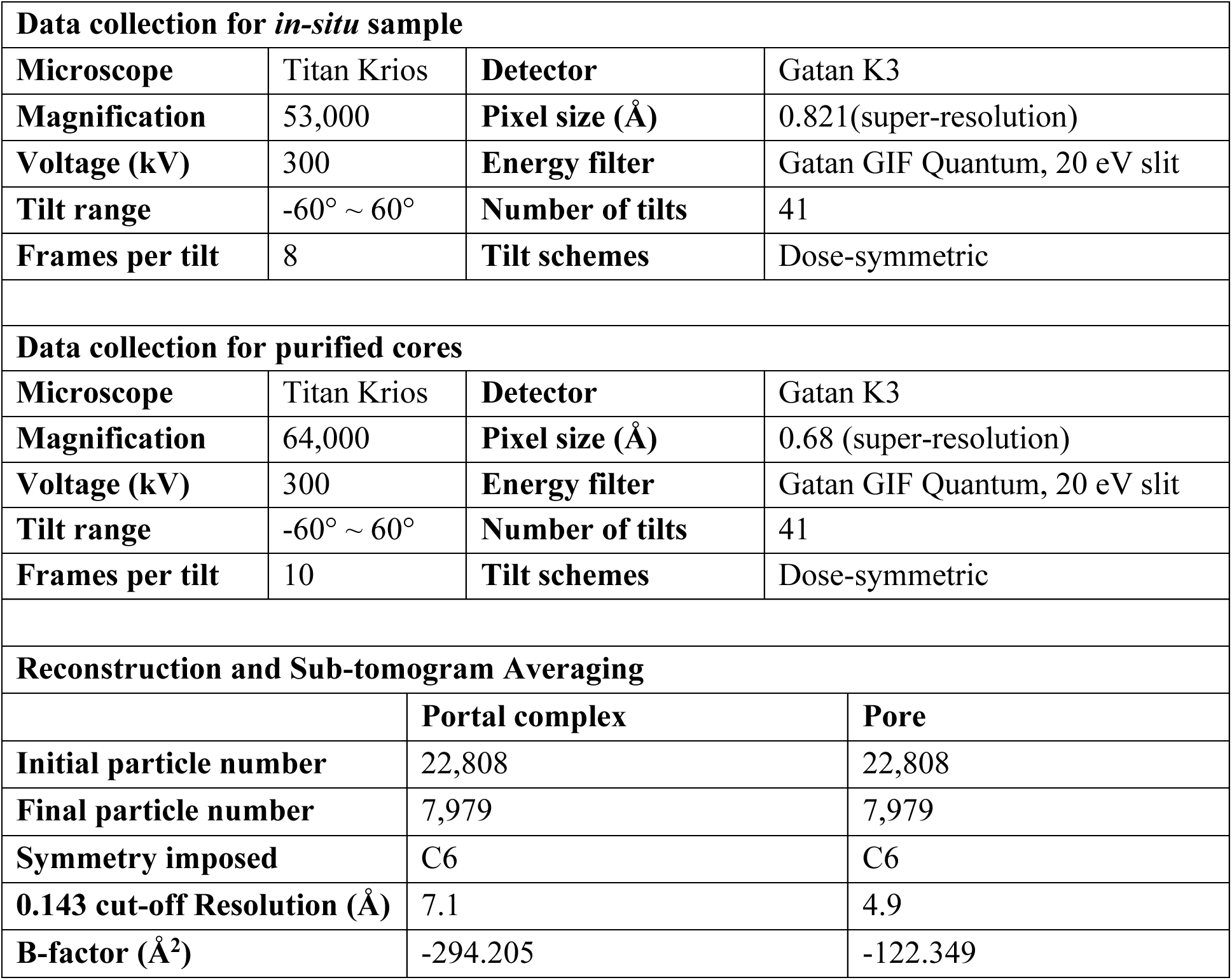
Cryo-ET data collection and STA parameters.

**Supplementary Table 2.** Proteins interacting with E6.

| **Protein** | **Amino acid number** | **Interact with p4a<sup>11</sup>** | **Interact with A4<sup>11</sup>** | **Interact with p4b<sup>11</sup>** | **Collar-ring candidate** | **Skirt- & petticoat-ring candidate** |
| --- | --- | --- | --- | --- | --- | --- |
| **E6** | 567 | Yes | Yes | / |  |  |
| **ATI (A25)** | 725 | Yes | / | Yes |  | ✓ |
| **F8** | 65 | / | / | Yes |  |  |
| **I7** | 423 | / | / | / |  |  |
| **I6** | 382 | / | / | Yes |  | ✓ |
| **RAP94 (H4)** | 795 | Yes | / | / |  |  |
| **VPK2 (F10)** | 439 | / | Yes | / |  |  |
| **M1** | 472 | Yes | / | Yes |  | ✓ |
| **F17** | 101 | Yes | Yes | Yes |  |  |
| **NPH2 (I8)** | 676 | Yes | Yes | / |  |  |
| **PAP1 (E1)** | 479 | Yes | / | / |  |  |
| **A12** | 192 | Yes | Yes | Yes |  |  |
| **RP19 (A5)** | 164 | / | / | Yes |  |  |
| **TOP1 (H6)** | 314 | Yes | / | / | ✓ |  |
| **DUSP (H1)** | 171 | / | / | / |  |  |
| **MCE (J3)** | 333 | Yes | / | / | ✓ |  |
| **L3** | 350 | / | Yes | Yes |  | ✓ |
| **ETF1 (D6)** | 637 | Yes | / | Yes |  | ✓ |
| **E8** | 273 | / | / | / | ✓ |  |
| **A25** | 76 | / | / | / |  |  |
| **DNLI (A50)** | 552 | Yes | / | Yes |  | ✓ |
| **A27** | Attaches to viral membrane and locates on virus flat surface<sup>12,13</sup> |  |  |  |  |  |
| **G9** | Entry-fusion complex protein, locates on small tip of the virus<sup>12</sup> |  |  |  |  |  |
| **P4a and A4** | Palisade protein<sup>1-3</sup> |  |  |  |  |  |

**Supplementary Table 3.** Model refinement and validation.

| **Composition** |  | 596 |
| --- | --- | --- |
| Number of chains |  | 24 |
| Number of protein residues |  | 8,304 |
| Number of atoms |  | 68,388 |
| ligands |  | none |
| **R.M.S deviations** |  |  |
| Bond length |  | 0.004 Å |
| Bond angle |  | 0.948° |
| **Ramachandran plot** |  |  |
| Favored |  | 83.13% |
| Allowed |  | 15.86% |
| Disallowed |  | 1.02% |
| **Validation** |  |  |
| MolProbity score |  | 2.50 |
| Clash score |  | 20.55 |
| Poor rotamers |  | 0.16% |

